# Alteration of nociceptive Schwann cells in a mouse model of high-fat diet induced diabetic peripheral neuropathy

**DOI:** 10.1101/2024.03.12.584541

**Authors:** Guillaume Rastoldo, Ali K. Jaafar, Aurélie-Paulo-Ramos, Katy Thouvenot, Cynthia Planesse, Matthieu Bringart, Marie-Paule Gonthier, Gilles Lambert, Steeve Bourane

## Abstract

Diabetic peripheral neuropathy (DPN) is characterized by progressive and symmetrical sensory abnormalities and constitutes one of the earliest and main complications of diabetes. DPN is characterized by heterogeneous sensory symptoms such as chronic pain, tingling, burning or loss of sensation. Nociceptive Schwann cells (nSCs), a recently identified subtypes of dermal Schwann cells support terminal nerve fibers in mouse skin and contribute to mechanical sensation and neuropathic pain. While terminal nerve fibers density is evaluated in DPN models, there is currently no data about nSCs number and integrity during the early stages of the disease. In the present study, we determined the quantitative differences in terminal nerve fiber density as well as nSCs number and cellular extensions between control and high-fat diet (HFD) induced diabetic mice presenting with a neuropathic phenotype. We also characterized L1CAM as a reliable and specific marker of nSCs, a cell type previously assessed by the expression of S100β and Sox10. Interestingly, we observed a decrease in intraepidermal nerve fiber density (IENFD) associated with a significant reduction of nSCs in the glabrous foot skin of neuropathic mice. Overall, this study identifies L1CAM as a new marker of nSCs and indicates that these cells are impacted in diabetic peripheral neuropathy.

## Introduction

One of the first and main complications of diabetes is diabetic peripheral neuropathy (DPN), affecting more than 60% of diabetic patients [1]. DPN involves damage to the peripheral nervous system, i.e., the sensory and motor nerves as well as the autonomic nervous system that control our organs. The most common symptoms of DPN are an impairment of sensitivity to pain and touch [2]. The early stages of the pathology are characterized by abnormal sensory symptoms such as pain, tingling, burning sensations, electrical discharges, pain evoked by non-painful mechanical stimuli (allodynia), increased sensitivity to noxious stimuli (hyperalgesia) [2] significantly altering the patients’ quality of life. Other sensory deficits such as numbness are also observed with progressive loss of sensation in the lower limbs often leading to tissue damage with the development of ulcers, increasing the risk of amputation [3][4]. Reduced sensations in the foot also lead to progressive loss of balance resulting in an increased risk of falls [5, 6].

Despite decades of research, there is currently no effective treatment for DPN due to an incomplete understanding of the underlying molecular and cellular mechanisms. Much attention has been dedicated to the damage of small, non-myelinated nerve fibers (C-fibers) responsible for sensitivity to heat and pain. The degeneration and regeneration of these fibers are typical features of diabetes skin biopsy analyses serve as late-stage confirmations of DPN presence [7]. While C-fibers were traditionally believed to terminate freely in the epidermis, recent work by Abdo et al. challenged this concept [8]. Their findings reveal a close connection between C-fibers and specialized nociceptive Schwann cells (nSC) capable of independently generating pain signals. These cells are currently identified by the expression of S100β and Sox10 [9–11] and most importantly, eliminating nSC alone is sufficient to induce neuropathic pain in mice [11]. Considering these recent breakthroughs, we hypothesize that neuropathies involving C-fiber degeneration, as seen in DPN, may result from damages to nSCs, as suggested by a recent report demonstrating a decrease in nSC density in Streptozotocin-induced type 1 diabetic mice [9]. To date, no study has evaluated the integrity of nSC in a type 2 diabetes model. We have therefore explored the impact of type 2 diabetes on the integrity of nSC during DPN pathology development.

## Methods

### Research Design

All reported experiments were performed at the GIP-CYROI platform’s animal facility, conducted in accordance with the French and European Community Guidelines for the Use of Animals in Research (86/609/EEC and 2010/63/EU), and were approved by the local Ethics Committee (n°114) for animal experimentation and the French authorities (APAFIS#31763-2018020912188806, approved on May 21, 2021).

### Animals

Twenty C57BL/6J mice (male, 6 weeks old) were purchased from Janvier (Le Genest Saint Isle, France) and housed (5 mice per cage) in a temperature-controlled environment with a 12–12h light/dark photocycle. After 2 weeks of adaptation, mice were divided in two groups (n=10/group) and placed either on a control diet (10% kcal %fat; SF13-081) or a high-fat diet (45% kcal %fat; SF04-001) from Specialty Feeds during 16 weeks. Mice with fasting blood glucose above ≥ 200 mg/dL were considered diabetic [12].

Mice expressing a tamoxifen-inducible Cre recombinase under the control of the PLP promoter (PLP-CreER^T2^; [13]) were crossed to Ai14 mice, which have a LoxP-flanked stop cassette excisable by Cre. At 6 weeks of age, the PLP-CreER^T2^; Ai14 progeny were injected intraperitoneally with tamoxifen (100 µg/g of mice) for 5 consecutive days. Tamoxifen was prepared in a mixture of 9/10 corn oil and 1/10 ethanol, and stored at 4 °C in a light-proof container. After 2 weeks, the mice were euthanized and the tissue collected for immunostaining.

### Genotyping

Genotyping of progeny from PLP-CreER^T2^ and Ai14 mice were done with PCR using CreDN2 (GAT CTC CGG TAT TGA AAC TCC AGC), CreUP2 (GCT AAA CAT GCT TCA TCG TCG G), Ai14mut-Fwd (CTG TTC CTG TAC GGC ATG G) and Ai14mut-Rev (GGC ATT AAA GCA GCG TAT CC) primers.

### Oral glucose tolerance test (OGTT)

An OGTT was performed after 16 weeks of diet in all mice. After 6 hours of fasting, one drop of tail blood was collected and glycemia was analyzed using a standard glucometer (One Touch Profile, Lifescan Inc., Milpitas, CA, USA, #6 strips). Oral gavage with glucose 30% (2g/kg body weight) into conscious mice was performed and glycemic levels measured 15, 30, 45, 60, 90 and 120 min post-administration.

### Nerve conduction velocity

Mice were anesthetized by using isoflurane (IsoFlo^®^, Centravet, France) and placed on a heating pad. Once anesthetized, motor nerve conduction velocities (MNCV) and sensory nerve conduction velocities (SNCV) were determined as previously described [14–16]. Using the Keypoint 4, Dantec-Medtronic electroneuromyography device, we performed both proximal and distal nerve stimulations of the sciatic nerve, registering the transmitted potentials with sensing electrodes at the gastrocnemius muscle. We used stainless steel subdermal needle electrodes to deliver single square-wave supramaximal stimulation with 0.2 millisecond impulses. MNCV was calculated by subtracting the distal from the proximal latency (measured in milliseconds) from the stimulus artifact of the take-off of the evoked potential, and the difference was divided into the distance between both stimulating electrodes (measured in millimeters using a Vernier caliper). For SNCV, recording electrodes were placed on the dorsum of the foot and stimulating electrodes on the ankle. The latency of onset (milliseconds) of the sensory nerve action potential after supramaximal antidromic stimulation of the sural nerve at the ankle was divided by the distance between the recording and stimulation electrodes (measured in millimeters using a Vernier caliper).

### Tissue Harvest

After 16 weeks of diet, mice received a subcutaneous injection of Buprenorphine (0.05 mg/kg, Buprecare^®^, Centravet, France) and were anesthetized by using isoflurane (3%). Mice were then euthanized by intracardiac puncture and blood samples were collected and centrifuged at 2000 rpm for 10 min at 4°C. Serum was collected and stored at −80°C until analysis. Intracardiac perfusion was performed with 20 mL of phosphate buffered saline (PBS) and 20 mL of 4% paraformaldehyde (PFA) dissolved in PBS. The hind-paw plantar surfaces of each mouse were collected and post-fixed overnight in 4% PFA solution. Biopsies of foot skin were then rinsed and cryoprotected into 30% sucrose dissolved in PBS at 4°C. The samples were then embedded in optimal cutting temperature (OCT, Tissue-Tek^®^, USA) compound and stored at −80°C until sectioning.

### Immunohistochemistry

Tissus sections (30 μm) were collected from the hind-paw plantar surface and washed (3x10min) with PBS. Blocking and permeabilization were done by incubation (1h) in 5% donkey anti-serum and 0.3% Triton X-100. For intraepidermal nerve fibers density, sections were incubated with the primary antibody anti-PGP9.5 (1:500, #14730-1-AP, ProteinTech) for 2 days; first overnight at 4°C and the next day sections were incubated overnight at room temperature. The same protocol was applied for L1 cell adhesion molecule (L1CAM, 1:500, #20659-1-AP, ProteinTech) antibody but the sections were incubated only overnight at 4°C. Sections were then rinsed in PBT (PBS + 0.1% triton) and incubated for 1 h at room temperature with the secondary antibody Alexa Fluor 594 donkey anti-rabbit (1:500, Abcam; ab150064), or Alexa Fluor 488 donkey anti-Rabbit (1:500, Invitrogen; A-21206).

### Identification and quantification of nociceptive Schwann cells

Nociceptive Schwann cells were identified according to Hu et al. criteria [9]. Commonly, nSCs connect to PGP9.5+ nerve endings and are located within 25 µm depth in the subepidermal zone at the junction between dermis and epidermis. For quantification, we excluded L1CAM positive staining in glands or large peripheral fiber bundles found deeper in the dermis. The number of nSCs was reported per mm length of foot skin. To quantify nSCs cellular extensions, we measured cell fluorescence using ImageJ. We selected the region of interest (ROI) at the subepidermal zone in each image of the foot skin and measured the mean fluorescence value in that area using ‘Mean gray value’.

### Confocal imaging

Intraepidermal nerve fibers density and nociceptive Schwann cells number were evaluated under a laser scanning confocal microscope Eclipse (Nikon) with a 40x objective. Following immunostaining, 30 µm thick Z-stack images were taken and analysis was performed on orthogonal maximum intensity projection of the original confocal Z-stack images.

### Metabolic markers

Plasma hemoglobin A_1_C levels were determined using DCA^®^ Vantage Analyzer (Siemens Healthineers, France) and hemoglobin A_1_C reagent kit (#23-312018). Serum triglycerides (TG) and total cholesterol levels were determined using colorimetric assays (Diasys^®^, #157109910021 and #113009910021, Germany). Plasma insulin and leptin concentrations were measured using a mouse insulin and leptin ELISA kit (Mercodia^®^, #10-1247-01, Sweden and RayBiotech^®^, Peachtree Corners, GA, USA). All metabolic markers were measured after 16 weeks of diet.

### Behavioral tests

#### Von Frey test

Mice were familiarized with the testing apparatus 1h/day during a week before behavioral testing. Animals were placed on an elevated wire grid and the plantar surface of the hindpaw was stimulated with calibrated von Frey monofilaments (0.008-6 g). The paw withdrawal threshold for the von Frey assay was determined by SUDO up-down method [17]. Testing was performed every 4 weeks after the start of the diet and up to 16 weeks.

#### Hargreaves test

To measure radiant heat pain by Hargreaves test [18], animals were put in plastic boxes and the plantar paw surface was exposed to a beam of radiant heat according to the Hargreaves method. Paw withdrawal latency was then recorded and beam intensity was adjusted to result in a latency around 10 seconds in control animals. Testing began after exploration and after grooming behaviors ended. High speed of skin heating (6.5°C/sec) preferentially activates Aδ-fibers [19]. Based on previous study [20], we used a medium to high speed of temperature heating (5°C/sec during 10 sec to obtain 50°C) to predominantly activate Aδ-fibers. Testing began after exploration and grooming behaviors ended. Heat stimulation was repeated 5 times with an interval of 10 min for each animal and the mean was calculated. A cutoff time of 30 seconds was set to prevent tissue damage. Testing was performed every 4 weeks after the start of the diet and up to 16 weeks.

#### Brush Test

To measure dynamic allodynia, mice were familiarized with the testing apparatus (the same that for von-Frey test) 1h/day during a week before testing. A soft paintbrush (Lefranc & Bourgeois^®^ - Brush no. 4 #175160) was used to gently stroke the plantar surface of the hindpaw in the direction from heel to toe. The test was repeated five times, with intervals of 5 min. For each test, no evoked movement was scored as 0, and walking away or occasionally brief paw lifting (∼1 sec or less) was scored as 1. For each mouse, the cumulative scores of the five tests were used to form a percentage response (100% response equivalent to 5 evoked movements during the 5 stimulations of the paw). Testing was performed every 4 weeks after the start of the diet for up to 16 weeks.

#### Pinprick Test

To measure acute mechanical pain, mice were familiarized with the testing apparatus (the same that for von-Frey test) 1h/day during a week before testing. After this acclimation, for pinprick test, we touched the plantar surface of the hindpaw with a pin [21] and measured numbers of withdrawal response as for the brush tests. For each mouse, the cumulative scores of the five tests were used to form a percentage response (100% response equivalent to 5 evoked movements during the 5 stimulations of the paw). Testing was performed every 4 weeks after the start of the diet and up to 16 weeks.

### Statistical analysis

We first analyzed the normality of data and showed that the results of serum triglycerides and total cholesterol levels, insulinemia, motor NCV, sensory NCV, IENFD and L1CAM-positive cells were normally distributed. Thus, unpaired Student’s *t*-tests (two tailed) were used for statistical comparisons. For behavioral data, a two-way ANOVA for repeated measures followed by Bonferroni’s post-hoc test was used to determine statistically significant differences. A *P* value of less 0.05 was considered significant.

## Results

### 1. Body weight gain, serum triglycerides and total cholesterol levels

To determine the effect of high-fat diet (HFD) on the development of peripheral neuropathy, we fed mice with a control diet (10% kcal %fat) or a HFD (45% kcal %fat) for 16 weeks (n=10/group, **Figure 1A**). As anticipated, HFD-fed mice gained significant more weight compared with controls. The difference became significant from week 11 onward (**Figure 1B**). In line with weight gain, blood samples analyses revealed that HFD-fed mice exhibited higher circulating levels of leptin compared with controls (**Figure 1C**). Triglycerides levels were not significantly increased (**Figure 1D**). However, HFD-fed mice showed significantly higher total plasma cholesterol level than mice fed with the control diet (*P* < 0.01; **Figure 1E**).

**Figure 1:**
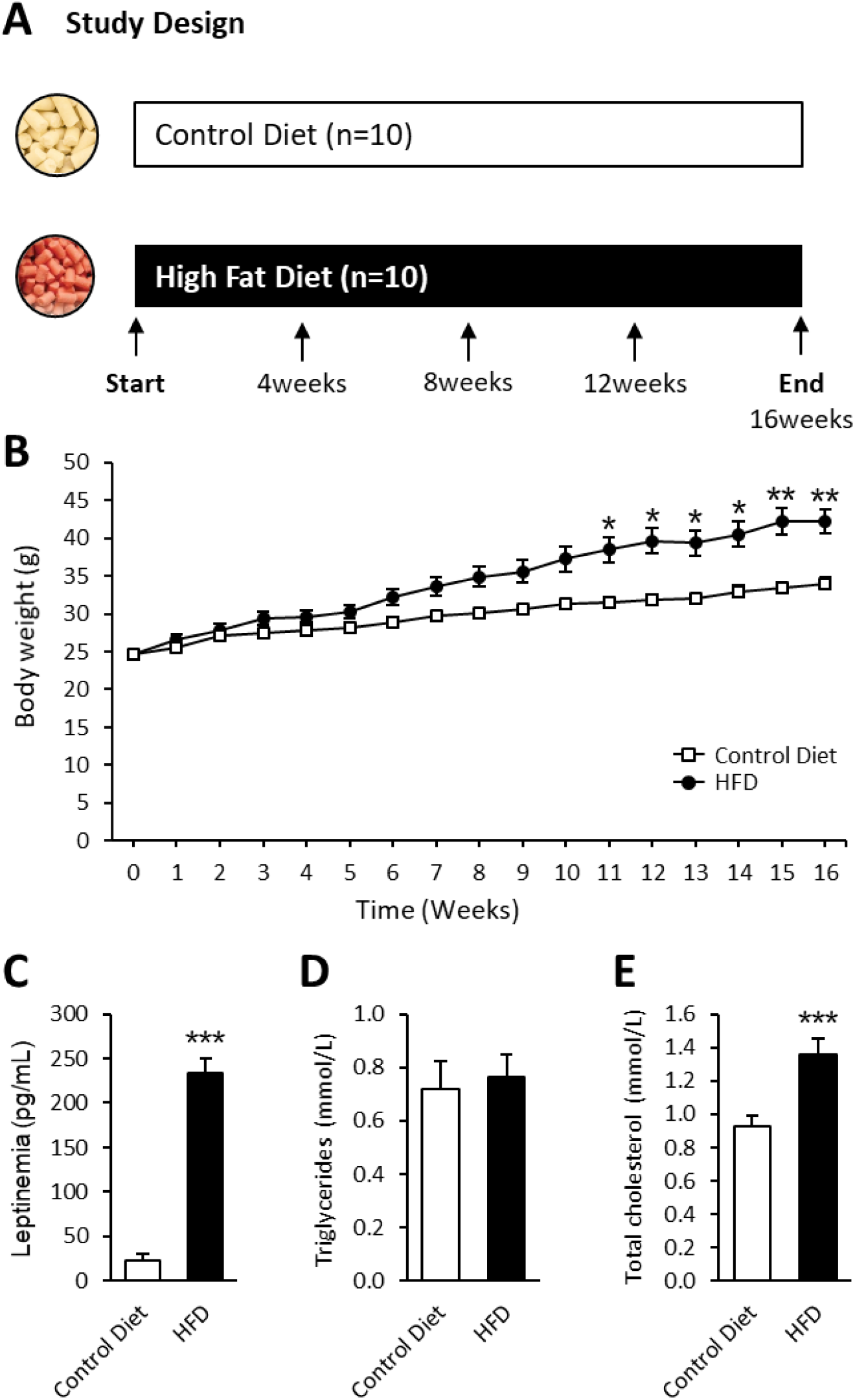
Study design and HFD-induced dyslipidemia. (**A**) Study Design used to assess the effect of a HFD (45% kcal %fat) vs. a control diet (10% kcal %fat) in two groups of mice (n=10/group). (**B**) Body weight of control diet-fed mice and HFD-fed mice during the 16 weeks of diet. (**C**) Leptinemia of control and HFD mice at 16 weeks of diet. (**D**) Plasma levels triglycerides of control and HFD mice at 16 weeks of diet. (**E**) Total cholesterolemia of control and HFD mice at 16 weeks of diet. \**P* < 0.05, \*\**P* < 0.01, \*\*\**P* < 0.001: control diet vs. HFD. Data are presented as means ± SEM.

### 2. Glucose homeostasis / Glucose regulation

Fasting glucose levels exhibited a significant increase in HFD-fed mice compared with controls (*P* < 0.001; **Figure 2A**) whereas HbA1c levels were not significantly altered (**Figure 2B**). In contrast, plasma insulin levels in HFD-fed mice were significantly increased compared with controls (*P* < 0.001; **Figure 2C**) indicating impaired glucose tolerance and insulin resistance. We performed an oral glucose tolerance test (OGTT) after 16 weeks of diet on both groups and observed that glucose tolerance was impaired (**Figure 2D**) with the AUC of blood glucose concentrations significantly higher in HFD-fed mice compared to control diet-fed mice (*P* < 0.01; **Figure 2E**).

**Figure 2:**
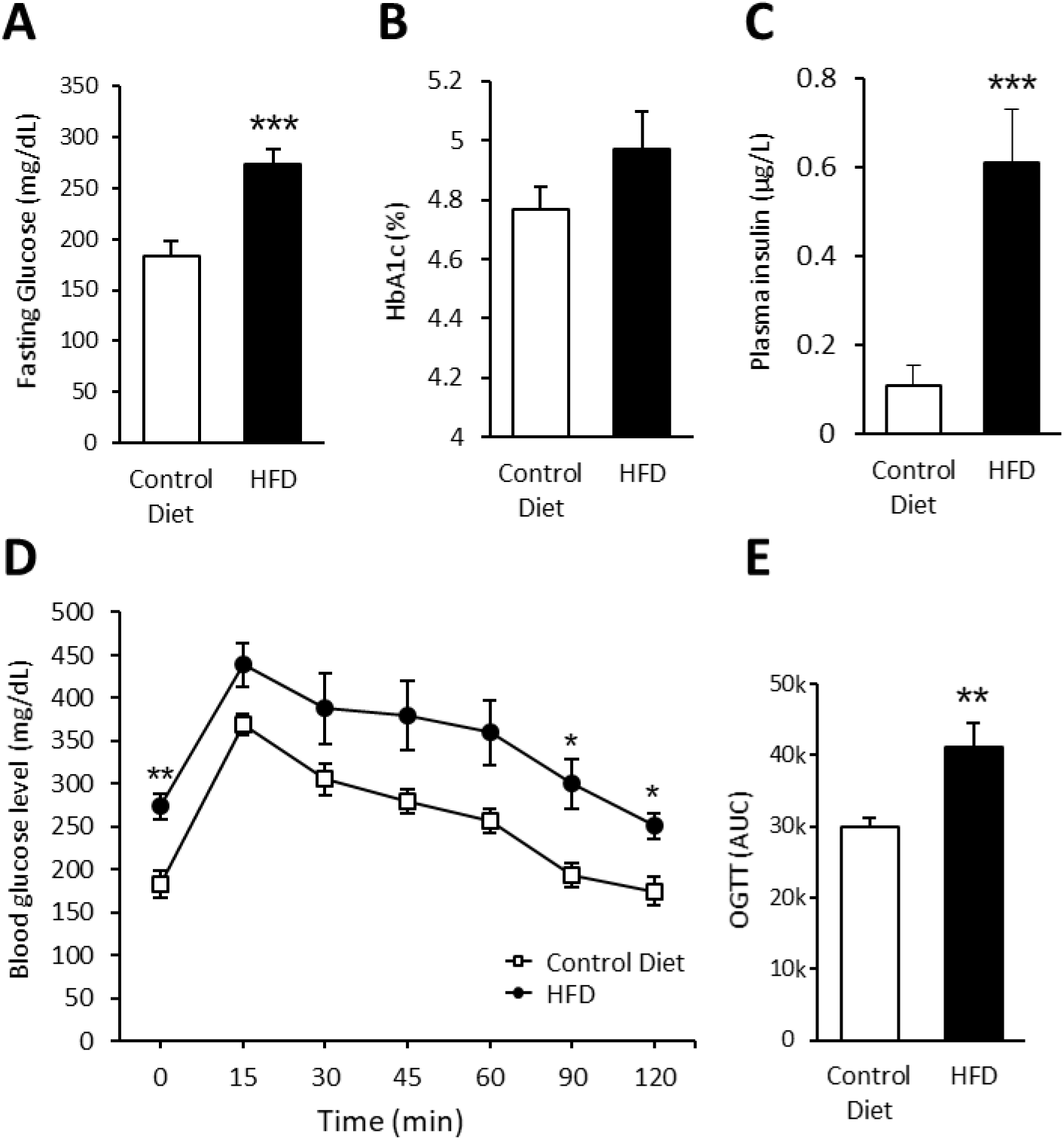
HFD-induced impaired glucose tolerance. **(A)** Fasting blood glucose of control diet-fed mice and HFD-fed mice at 16 weeks of diet. **(B)** Percentage of glycated hemoglobin (HbA1c) of control diet-fed mice and HFD-fed mice at 16 weeks of diet. **(C)** Plasma insulin of control diet-fed mice and HFD-fed mice at 16 weeks of diet. **(D)** Oral glucose tolerance test (OGTT) curves performed at 16 weeks of diet in control and HFD-fed mice. **(E)** Area under the curve (AUC) during OGTT for control diet-fed mice and HFD-fed mice. \**P* < 0.05, \*\**P* < 0.01, \*\*\**P* < 0.001: control diet vs. HFD. Data are presented as means ± SEM (n=10/group).

### 3. Nerve conduction studies

At week 16, nerve conduction velocity (NCV) was measured in the sciatic and sural nerves of each animal. HFD-fed mice did not show any significant change in motor (**Figure 3A**) or sensory (**Figure 3B**) NCV compared with controls.

**Figure 3:**
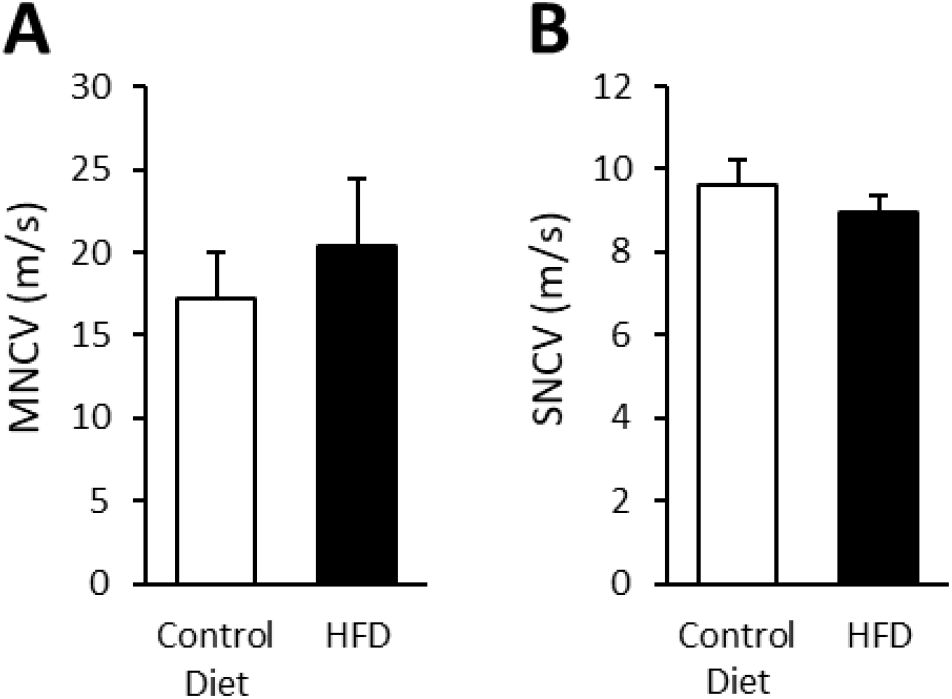
Nerve conduction studies. **(A)** Motor nerve conduction velocity (MNCV) performed at 16 weeks of diet in control and HFD-fed mice. **(B)** Sensory nerve conduction velocity (SNCV) performed at 16 weeks of diet in control and HFD-fed mice. Data are presented as means ± SEM (n=10/group).

### 4. Thermal and mechanical pain sensitivity

To evaluate peripheral neuropathy we performed various behavioral sensory tests every 4 weeks for 16 weeks. HFD-fed mice exhibited an increase in paw withdrawal threshold as early as 8 weeks (4 weeks HFD vs. 8 weeks HFD, *P* < 0.01, **Figure 4A**) and were significantly different from their control diet-fed counterparts after 12 weeks of diet (12 weeks control diet vs. 12 weeks HFD, *P* < 0.01, **Figure 4A**), indicating a decrease in mechanical sensitivity. In parallel, we observed an increase in thermal sensitivity of HFD-fed mice compared with control mice from week 8 (**Figure 4B**). The latency of hind-paw withdrawal in response to noxious thermal stimulus was significantly decreased in HFD-fed mice compared with control diet-fed mice at 12 and 16 weeks (*P* < 0.001). No significant change in haptic mechanical sensitivity (**Figure 4C**) evaluated with the brush test or acute mechanical pain sensitivity (**Figure 4D**) assessed by the pinprick test was noticed over 16 weeks between both groups.

**Figure 4:**
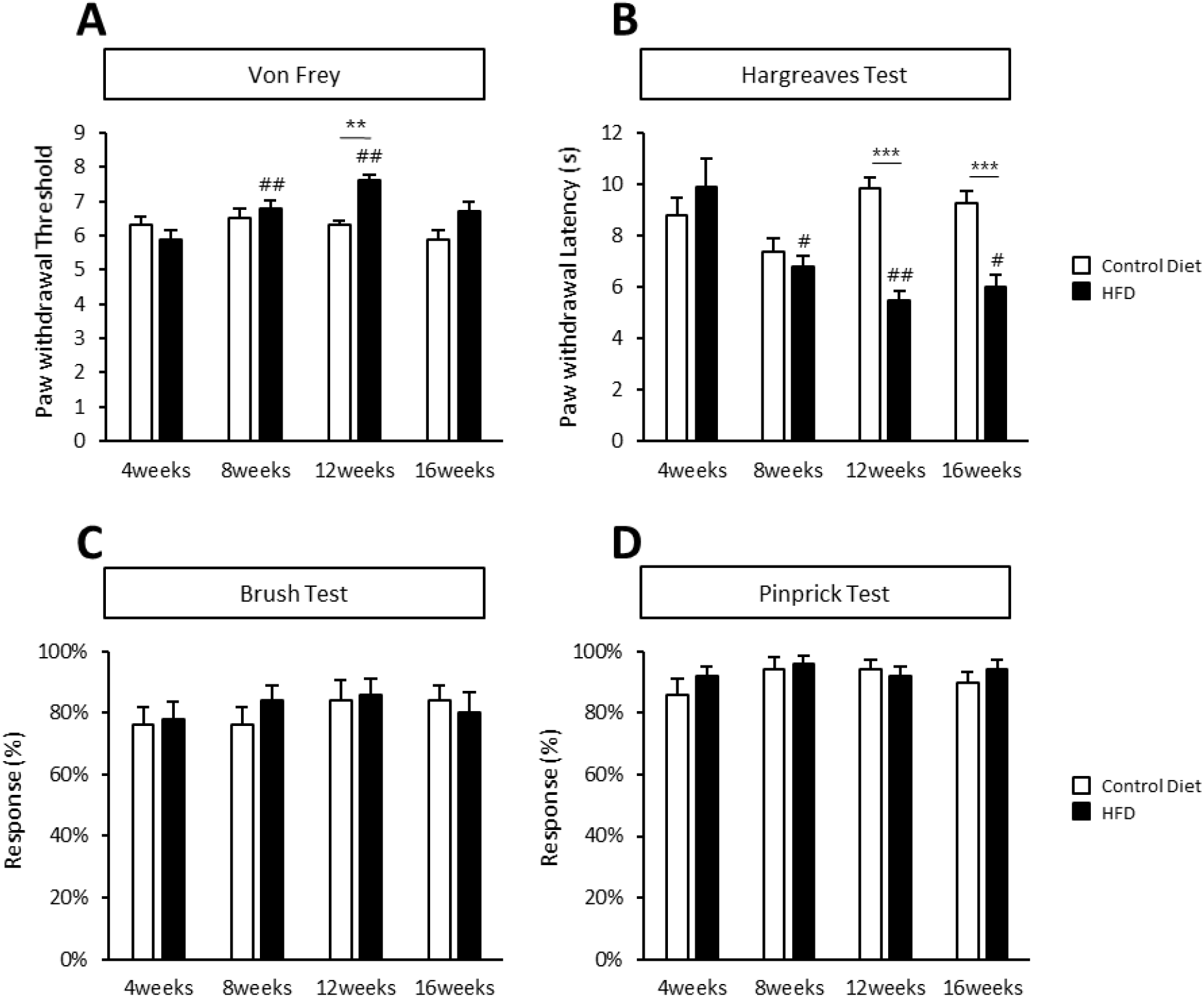
Mechanical and thermal sensitivity changes induced by HFD. **(A)** Quantification of static mechanical sensitivity was assessed every 4 weeks in control and HFD-fed mice using von Frey filaments. **(B)** Quantification of thermal sensitivity by Hargreaves test every 4 weeks in control and HFD-fed mice. **(C)** Quantification of haptic mechanical sensitivity by brush test every 4 weeks in control and HFD-fed mice. **(D)** Quantification of acute mechanical pain by pinprick test every 4 weeks in control- and HFD-fed mice. \*\**P* < 0.01, \*\*\**P* < 0.001: control diet vs. HFD. #*P* < 0.05, ##*P* < 0.01: HFD 4weeks vs. HFD 8, 12 or 16 weeks. Data are presented as means ±SEM (n=10/group).

### 5. Epidermal innervation and Nociceptive Schwann cells

To fully characterize the neuropathic phenotype in our model, we analyzed the intraepidermal nerve fiber density (IENFD) in the glabrous skin of control and HFD-fed mice. We found a significant reduction of foot skin innervation in HFD-fed compared with control mice after 16 weeks of diet (*P* < 0.001; **Figure 5A, 5B**). Additionally, we investigated the integrity of nociceptive Schwann cells that are present in the skin at the junction between the epidermis and the dermis by using immunohistochemistry against L1CAM that we found highly expressed in this cell type. To validate the specificity of L1CAM expression in nociceptive Schwann cells, we first verified its colocalization with Sox10 and S100β, two known markers of these cells [9–11] (**Figure 5C**). To further these observations, we confirmed the specific expression of L1CAM in td-Tomato positive nSCs in PLP-CreER^T2^;Ai14 double transgenic mouse foot skin (**Figure 5D**), [13, 22]. Altogether, these results indicate that L1CAM is a reliable and specific marker of nSCs.

**Figure 5:**
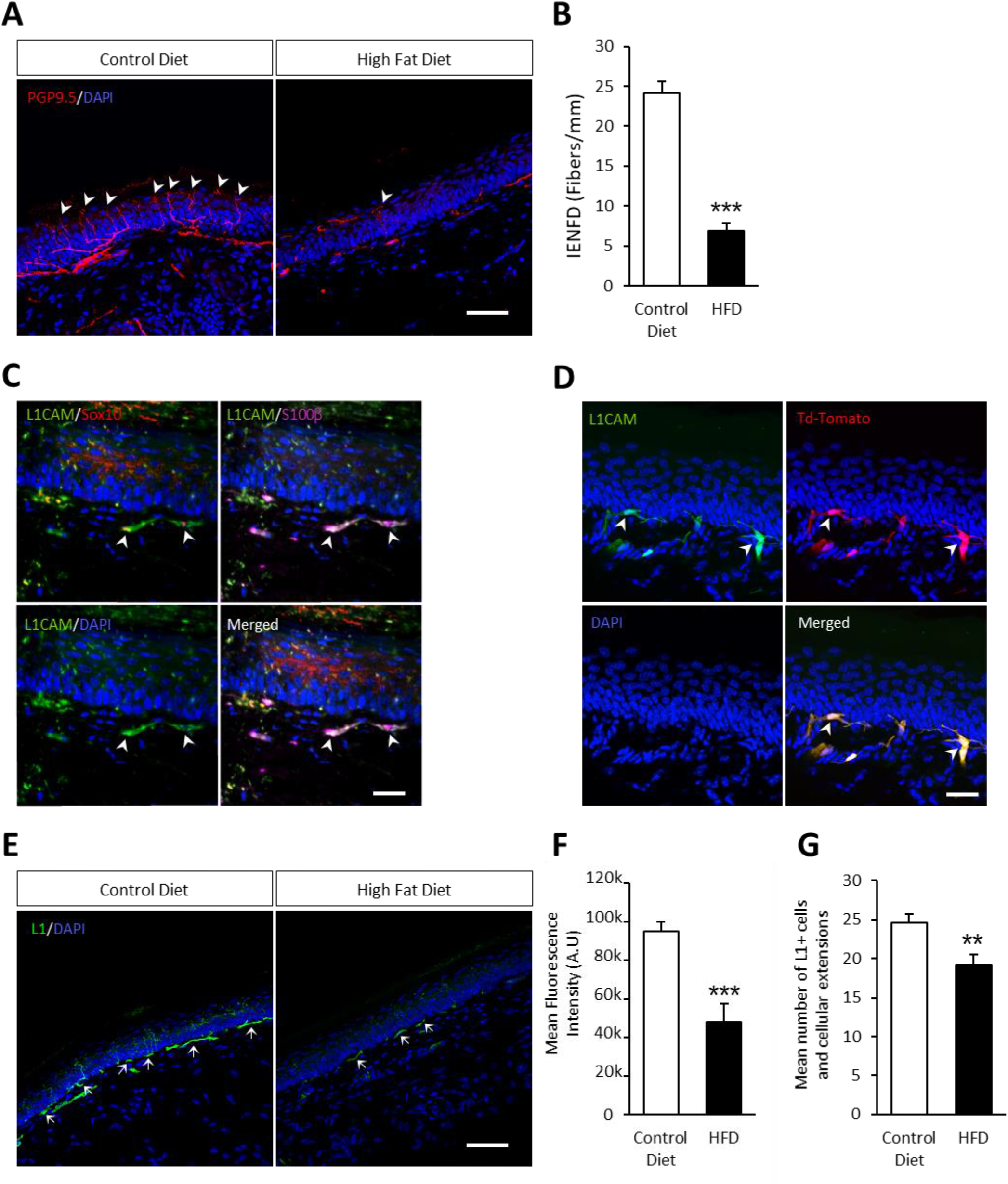
HFD-induced small nerve fiber and nociceptive Schwann cells degeneration in the foot skin. (**A**) PGP9.5 immunofluorescence with DAPI staining in foot skin section of control (n=5) and HFD-fed mice (n=5). Arrowhead indicate nerve fibers in the epidermis of the foot skin. (**B**) Quantification of IENFD is presented as the number of fibers/mm of epidermis. (**C**) L1CAM immunofluorescence colocalized with Sox10 andS100β classical markers of nociceptive Schwann cells in the foot skin. (**D**) L1CAM immunofluorescence colocalized with Td-tomato expressing nociceptive Schwann cells in a tamoxifen-induced PLP-CreERT2;R26TdTomato double transgenic mouse. Arrowhead indicate nociceptive Schwann cells and their cellular extensions at the border of the epidermis and dermis. (**E**) L1CAM immunofluorescence with DAPI staining in foot skin section of control (n=5) and HFD-fed mice (n=5). Arrows indicate nociceptive Schwann cells and their cellular extensions at the border of the epidermis and the dermis. **(F**) Mean fluorescence intensity quantification of L1CAM immunostaining at the localization of nociceptive Schwann cells (at the border of epidermis and dermis).(**G**) Quantification of L1CAM-positive cells and their cellular extensions presented in number/mm of epidermis. \*\**P* < 0.01, \*\*\**P* < 0.001: control diet vs. HFD. Data are presented as means ± SEM. Scale bar: 50 μm.

Using this new marker to identify nSCs, we observed that, both mean fluorescence intensity (**Figure 5E,F**) and mean number of L1CAM-positive cells and cellular extensions (**Figure 5E,G**) were significantly decreased in HFD-fed mice compared with control mice (*P* < 0.001 and *P* < 0.01 respectively). Altogether, these results indicate a decrease in both sensory nerve endings and number of nociceptive Schwann cells in the foot skin of HFD mice.

## Discussion

In the present study, we used HFD fed mice during 16 weeks to generate a mouse model of diabetic peripheral neuropathy and compared dyslipidemia, glucose tolerance, and sensory deficits of these mice with those on placed on a control diet. We observed that HFD-fed mice display increased weight gain, elevated leptinemia, hypercholesterolemia, impaired glucose tolerance, elevated fasting blood glucose and insulin resistance compared with control-fed animals, recapitulating the metabolic dysfunctions observed in type 2 diabetes. The analysis of IENFD of the foot skin revealed an important reduction in terminal nerve fibers and nSCs in HFD-fed mice compared with mice fed with a control-diet, paralleling behavioral sensory deficits in HFD-fed mice, with the development of mechanical hyposensitivity and thermal hypersensitivity, clearly indicative of a peripheral neuropathy phenotype.

To characterize the progression of peripheral neuropathy, we followed the guidelines of the diabetic neuropathy study group of the European Association for the Study of Diabetes. Namely, we assessed three key features present in human pathology (nocifensive behavior, nerve conduction velocity, and IENFD) considering that the presence of alterations in two of these three parameters is necessary to establish a neuropathic phenotype in rodents [23]. In this study, we demonstrate that HFD-fed mice exhibite mechanical hypoalgesia and thermal hyperalgesia coupled to a severe loss of intraepidermal nerve fibers but no impairment of motor or sensory nerve conduction velocity. According to the above-mentioned criteria of Neurodiab, we can assert that our HFD-fed mice developed diabetic peripheral neuropathy.

To identify possible damages to small sensory fibers, we next performed a quantification of intraepidermal nerve fibers density considered as the gold standard for the evaluation of small fiber neuropathies [24]. Damage to small sensory fibers is one of the earliest manifestations of DPN that can be observed in prediabetic conditions [25]. We found a 70% reduction in epidermal innervation in the hind paw of the HFD-fed mice in comparison with control-fed mice, in agreement with previous reports [26–28]. Studies have shown that small nerve fibers penetrating the epidermis are mostly nociceptive [7] and that these fibers undergo degeneration and regeneration at the early stages of type 2 diabetes in humans [5, 29]. Until recently, it was thought that because unmyelinated C-fibers lack the protection and nutrient supply that myelinated Schwann cells provide to A-fibers, C-fibers were more likely affected at the early stages of diabetic neuropathy. However, recently a specialized cutaneous type of Schwann cells, termed nociceptive Schwann cells, in close relationship with nociceptive C-fibers in the dermis has been discovered [8]. These cells contribute to terminal axon support and play a critical roles in skin mechano-sensitivity and neuropathic pain [10, 11, 34, 35]. Therefore, nociceptive endings in the epidermis are not free but form a mesh-like organ with a mutual dependence between C-fibers and nSCs. The absence of nSCs induces a retraction of epidermal nerve fibers and development of mechanical, cold, and heat hyperalgesia [11]. In the present study, we identified L1CAM as a reliable marker for nSCs, as an alternative to the conventional S100β / Sox10 staining methods [9–11]. As outlined by Hu et al., the precise localization of Schwann cells serves as a crucial criteria in distinguishing nSCs [9]. These distinctive cells are arranged within a 25 µm depth in the subepidermal region, excluding glands and large fiber bundles. Based on these criteria and the expression of L1CAM, we demonstrate a significant decrease in the number of nSCs and their cellular extensions in T2D mice, as previously observed in type 1 diabetic mice [9].

Hyperglycemia represents a key factor that can induce apoptosis in Schwann cells [37]. Inhibiting apoptosis of Schwann cells under high-glucose conditions is considered a promising approach to treat DPN [38]. However, glycemic control has little impact of the development and progression of neuropathy in type 2 diabetic patients [39]. Dyslipidemia is another frequent perturbation of T2D, but the impact of this lipids on Schwann cells integrity has received relatively little attention [40–42]. The elucidation of mechanisms implicated in the alteration of nSCs in HFD-fed mice therefore seems central for the future development of potential treatments strategies of DPN.

## Conclusion

In this study, we used HFD to induce a mouse model of diabetic peripheral neuropathy. We observed key metabolic dysfunctions indicative of T2D in these animals. Behavioral analysis uncovered early-onset of peripheral neuropathy with mechanical hypoalgesia and thermal hyperalgesia. We also identified L1CAM as a new marker of nociceptive Schwann cells. Interestingly, we observed a significant decrease in number of nociceptive Schwann cells and their extensions in the foot skin of neuropathic mice, highlighting the vulnerability of these cells in T2D conditions. Overall, our analyses recapitulate the early events in diabetic peripheral neuropathy pathophysiology, paving the way for future investigations into the molecular mechanisms underlying nociceptive Schwann cells alterations and potential strategies to thwart this debilitating complication of diabetes.

## Acknowledgment

We thank Dr Douglas Wright and Janelle Ryals for their assistance with the PGP9.5 immunohistochemistry. We thank Dr Aziz Moqrich for providing the PLPcreERT2 and Ai14 mice.

## Data availability

The authors confirm that the data supporting the findings of this study are available within the article and its supplementary materials. Further information and requests may be sent to the corresponding author.

## Funding

This research was supported by a grant funded by Inserm, Paris, France [ATIP-AVENIR].

## Contribution statement

All authors meet the ICMJE uniform requirements for authorship and approved the final version of the manuscript. GR and SB conceptualized the study and contributed to study design. GR, AKJ, AP and SB participated in skin biopsy and data acquisition. MPG and KT performed plasma insulin and leptin concentrations measurements. GR and MB participated in OGTT. GR and AKJ participated in data analysis. MB and CP took care of the maintenance of the animals. GL and MPG helped in critical reading of the article. All authors contributed to manuscript drafting, editing and revision. GR and SB wrote the original draft and all authors revised the manuscript. SB accepts full responsibility for the work and/or the conduct of the study, had access to the data, and controlled the decision to publish.

## Notes

### Competing Interest Statement

The authors have declared no competing interest.

### Summary of Updates

This version of the manuscript has been revised to update the results and the discussion part.

